# Pseudogenization, genome streamlining and specific gene repertoire landmark the genomes of *Carnobacterium maltaromaticum* isolated from diseased sharks

**DOI:** 10.1101/600684

**Authors:** Laura Martinez Steele, Christopher G Lowe, Mark S Okihiro, Jesse G. Dillon, Renaud Berlemont

## Abstract

*Carnobacterium maltaromaticum* is a well-known pathogen of bony fish. More recently, *C. maltaromaticum* have been isolated from the brain and inner ear of disorientated and stranded common thresher (*Alopias vulpinus*) and salmon shark (*Lamna ditropis*). While thresher shark strandings are recent, salmon sharks have been stranding for decades, suggesting a long-term association between *C. maltaromaticum* and sharks. Interestingly, some strains of *C. maltaromaticum* are used by the food industry for their probiotic and antimicrobial activity. Here, we sequenced the genome of 9 *C. maltaromaticum* strains (SK-isolates) from diseased common thresher and salmon sharks and compared them to other *C. maltaromaticum* strains in order to identify the genomic signatures that differentiate the disease-associated from the innocuous *C. maltaromaticum* isolates. SK strains formed a monophyletic clade, with a conserved gene repertoire, and shared a high degree of pseudogenization even though isolates were from different shark species, locations, and across years. In addition, these strains displayed few virulence associated genes and unique genomic regions, some resulting from horizontal gene transfer. The association of diseased sharks and SK strains suggests their role as potential pathogens. Although the high degree of pseudogenization suggests a transition to a host-adapted lifestyle, a set of conserved functional genes highlights the need of essential functions required for a host-independent life style. Globally, this work identifies specific genomic signatures of *C. maltaromaticum* strains isolated from infected sharks, provides the framework to elucidate the role of SK strains in the development of the disease in sharks, and further investigate the dissemination of SK strains in populations of wild fish.

## Introduction

Occurrences of stranded salmon sharks (*Lamna ditropis*) [1] and common thresher sharks (*Alopias vulpinus)* [2] have been happening along the West Coast of North America. These stranding events involve disoriented, sick juveniles that swim onto the beach, and occur throughout the year although mostly reported from July to September [3]. Both thresher and salmon sharks belong to the order of Lamniformes, and are top predators that migrate between the coastal and offshore waters of the Northeast Pacific Ocean, and use the continental shelf as nurseries when juveniles [4]. Histology of the brain and inner ear of stranded sharks revealed severe meningitis [1] and acute otitis, most probably caused by *Carnobacterium maltaromaticum* (phylum Firmicutes) identified in bacterial cultures obtained from damaged tissues [1, 2]. *C. maltaromaticum* is a facultative anaerobic, psychrotrophic, lactic acid bacterium, of biotechnological interest due to its antimicrobial properties inhibiting *Listeria monocytogenes* [5, 6]. It is also detected in many natural environments including the gastrointestinal tract of many teleost [7–11], where it is thought to stimulate the immune system, and thus is sometimes used as a probiotic in aquaculture [12, 13]. However, *C. maltaromaticum* is also a pathogen in cold-water teleost fishes exposed to various stressors (e.g., spawning, handling events) [11, 14]. In diseased fish, *C. maltaromaticum* is associated with pseudokidney disease, septicemia, splenomegaly, internal hemorrhages, muscular abscesses, visceral congestion, and thickening of the swim-bladder walls [11, 14, 15].

As of June 2018, the genus *Carnobacterium* comprises 11 species and 42 genomes are publicly accessible through the NCBI database (S1 Table, S2 Table). Genomes range from 1.5 to 4.0 Mbp and have GC content ranging from 34 to 39.4%. Among sequenced genomes, *C. maltaromaticum* strains have the largest, yet variable, genomes (3.3 ± 0.7 Mbp) mirroring the gain and loss of massive DNA fragments [16]. Additionally, *C. maltaromaticum* is the only species to be reported in multiple environments including processed food, human skin, and the digestive system of some teleost species [6].

Here, we characterized the genome of 9 *C. maltaromaticum* (SK strains) isolated from the brain and inner ear of stranded diseased-sharks (i.e., common thresher shark and salmon shark) from along the coast of North America between 2013 and 2016. Despite varying sample origin and collection times, we hypothesized that these strains would be phylogenetically related and would display specific genomic signatures discriminating them from strains originating from other sources. Thus, we first investigated the phylogenetic placement of the newly sequenced strains relative to publicly accessible *Carnobacterium* genomes using 16S rRNA sequence, tetranucleotide correlation search (TCS), and based on single nucleotide polymorphisms (SNPs) distribution in the core genome of all sequenced *C. maltaromaticum.* Next, we compared the functional potential, based on annotated genes, in the nine SK strains and in all the sequenced *C. maltaromaticum* (n_total_=17) in relation to their known habitat and lifestyle (i.e., food-derived and diseased teleost or sharks).

The consistent identification of the SK strains in stranded sharks using different techniques such as cytology, histology and bacterial culture [1, 2], suggests the potential role of *C. maltaromaticum* in the sharks’ disorientation and stranding. Thus, we specifically investigated the distribution of potential genes involved in pathogenicity. Furthermore, we searched for differences in the proportion of functional genes discriminating diseased-shark isolates and food-associated strains, to identify niche-specific functions associated with the origin of the strains [17, 18]. We also identified regions of lateral transfer such as “Genomic Islands” (GIs) [19, 20] and bacteriophages [21–23]. Finally, we determined the accumulation of pseudogenes [24, 25] in SK strains, relative to the other strains, as an indication of the reductive evolutionary processes often observed in pathogens [26–28].

For the first time, this study investigates the genomic signatures of *C. maltaromaticum* isolated from diseased-sharks in comparison to diseased-teleost and food relatives with the purpose of elucidating evolutionary forces that shaped the genome of strains associated with disease in wild-fish populations.

## Materials and methods

### Strains

Nine strains of *C. maltaromaticum* (SK strains) were isolated from the brain and inner ear of stranded common thresher sharks and salmon sharks and sequenced as described before (S1 Table) [2]. For phylogenetic placement, genomic DNA extracts were PCR-amplified using the primers GM3/GM4 [29] targeting the 16S rRNA gene and the products were Sanger-sequenced and deposited to the NCBI (S1 Table). Sequences were BLASTed against the NCBI database and were confirmed as *Carnobacterium maltaromaticum* (>99% identity).

For the comparative genome analysis, we first used 41 publicly accessible genomes from NCBI database covering different species from the genus *Carnobacterium*, and five pathogenic outgroups (S2 Table). Then we characterized 17 *C. maltaromaticum* genomes including 9 SK strain genomes (Table 1) and 8 isolated from various environments. The strain 757 CMAL (human isolate) was excluded from the *C. maltaromaticum* analysis because of its divergence from the rest of the strains, which might potentially interfere with the analysis. For consistency, contigs less than 3 kb were removed from all 46 genomes and were consecutively (re)annotated with the Rapid Annotation using Subsystem Technology (RAST) online server (v 2.0) [30, 31].

**Table 1.** Analyzed *C. maltaromaticum* genomes.

| Strain | Size (Mbp) | CDS | GC (%) | Contigs | Contigs > 3 Kb | N50 | Pseudo genes | Source/ Life-style | Ref. |
| --- | --- | --- | --- | --- | --- | --- | --- | --- | --- |
| **3-18** | 3.5 | 3245 | 34.3 | 160 | 71 | 82,960 | 41 | Pork | [32] |
| **ATCC 35586** | 3.5 | 3333 | 34.5 | 74 | 63 | 106,236 | 70 | Diseased trout | [33] |
| **DSM 20342** | 3.7 | 3481 | 34.3 | 132 | 54 | 79,358 | 48 | Milk with malty flavor | [34] |
| **DSM 20342 MX5** | 3.9 | 3636 | 34.4 | 5 | 5 | 3,620,667 | 71 | Milk with malty flavor | [34] |
| **DSM 20722** | 3.6 | 3301 | 34.3 | 60 | 22 | 133,412 | 41 | Vacuumed meat | [35] |
| **DSM 20730** | 3.5 | 3254 | 34.4 | 143 | 28 | 75,250 | 19 | Diseased trout | [35] |
| **LMA 28** | 3.7 | 3472 | 34.5 | 1 | 1 | NA | 155 | Cheese | [36] |
| **ML 1 97** | 3.2 | 3108 | 34.4 | 229 | 176 | 26,642 | 283 | Fresh salmon | [32] |
| **SK AV1** | 3.3 | 3220 | 34.3 | 42 | 18 | 312,952 | 262 | Thresher shark 1 brain | [2]<br>This study |
| **SK AV2** | 3.3 | 3219 | 34.3 | 123 | 19 | 312,934 | 264 | Thresher shark 1 ear |  |
| **SK AV3** | 3.3 | 3222 | 34.3 | 44 | 18 | 312,857 | 263 | Thresher shark 8 brain |  |
| **SK AV4** | 3.3 | 3220 | 34.3 | 44 | 18 | 312,883 | 262 | Thresher shark 8 ear |  |
| **SK AV5** | 3.4 | 3300 | 34.4 | 3,070 | 33 | 312,904 | 263 | Thresher shark 11 brain |  |
| **SK AV6** | 3.3 | 3225 | 34.3 | 116 | 19 | 312,881 | 263 | Thresher shark 11 ear |  |
| **SK LD1** | 3.3 | 3220 | 34.3 | 70 | 18 | 389,499 | 262 | Salmon shark 1 Brain |  |
| **SK LD2** | 3.3 | 3218 | 34.3 | 79 | 17 | 312,874 | 265 | Salmon shark 1 ear |  |
| **SK LD3** | 3.3 | 3219 | 34.3 | 76 | 19 | 312,880 | 263 | Salmon shark 2 brain |  |

Genome characteristics are extracted from the RAST except for the pseudogenes, which were
predicted using DFAST, and contigs which were taken from the NCBI.
Number of contigs are shown before and after contigs < 3 Kb were removed, while the rest of the
features shown are after contigs < 3 Kb were removed

### Phylogenetic placement

The nine SK strains were phylogenetically identified by comparing the complete 16S rRNA gene sequences from 28 *Carnobacterium* genomes with entire 16S rRNA sequence and 5 pathogenic outgroups (n_total_= 42 strains, S2 Table). Sequences were aligned using ClustalW [37] and we constructed a phylogenetic tree using the PHYLIP-phylogeny interference package (version 3.695, F84 distance method, and neighbor-joining tree) [38]. The tree was visualized using the interactive tree of life (iTOL v4) [39].

### Phylogenomic placement

We first performed TCS [40] against the JSpeciesWS database [41], and Average Nucleotide Identity (ANIb) among the newly sequenced genomes [42]. Next, we used the Pangenome Analysis Tool from Panseq [43] to identify SNPs in the core genome of the 17 *C. maltaromaticum* species only. We used default identity cut-off of 85% to exclude genes acquired recently through horizontal gene transfer. Next, a tree based on identified SNPs was built as previously described in the phylogenetic analysis of this study. Finally, we computed a clustering analysis of the genomes based on their functional distribution (FigFam-based annotation [30, 31]), using the pairwise Bray-Curtis dissimilarity index and visualized using complete linkage clustering [44] (n_total_= 46 genomes, S2 Table).

### Orthologous clustering and pan-genome functional annotation

Anvi’o pangenomic workflow [45] was used to identify and annotate gene clusters with the 2014 COG database [46]. Briefly similarities between genes were performed using BLASTp and weak matches were eliminated with *minibit heuristic* [45]. Next, clustering was performed with the Markov Cluster algorithm (MCL) [47] using default parameters with the exception for a mcl-inflation of 10 (used for closely related strains). The core, accessory and unique genes and their COG function and category distribution were extracted and used for further analysis. For this study core genome was defined as the number of genes present in all genomes, while accessory genes were those gene clusters present in more than 1 but less than 17 genomes, and finally singletons were genes clusters just found in one genome.

Because changes in functional distribution mirrors adaptation to different environments, we performed Wilcoxson Rank Sum test with Bonferroni correction for multiple comparison to detect differences in the COG categorical functions between isolates from food (n=6), diseased trout (n=2), and diseased shark (n=9).

### Pan-genome profile

A panmatrix containing gene clusters presence in 17 *C. maltaromaticum* genomes was imported from Anvi’o [45] to the Micropan package implemented in R [48] to generate pan-genome visualizations [49]. The Heaps law model [50] and the Binomial Mixture Model, using the Bayesian Information Criterion (BIC) [51], implemented in the package were used to determined openness of the pan-genome and to estimate pan and core-genome size when envisioning an “infinite” number of genomes. Rarefaction curve using 100 permutations helped visualize the increase of the pan-genome size with the addition of new genomes.

### Virulent associated genes

We searched for virulence factors, within the 17 *C. maltaromaticum* genomes, using protein sequences provided by the RAST annotation, and by performing a BLAST search (60% coverage and 60% identity) against the Virulence Factor Database (VFDB, [52]). These results were combined with genes under the *Virulence, Disease and Defense* category from the corresponding FigFam-annotated genomes [30, 31].

### Genomic plasticity

GIs were identified and visualized with Islandviewer v.4 [53] using annotation files retrieved from the RAST. *C. maltaromaticum* LMA28 was used as a reference to re-order contigs in draft genomes with default parameters. To determine unique islands in the SK strains we compared them to islands detected in the other strains using BLAST (60% coverage and 60% identity). Prophage sequences were predicted in re-ordered genomes using PHASTER [54]. This tool scores the results according the completeness of the sequence taking into account length, gene content, GC content and attachment sites. Results were labeled as intact, questionable and incomplete. Intact phage sequences were than scan for virulence genes as previously explained. The script “rod_finder” [55] was used to determine regions of difference (RODs) in each genome in order to determine DNA sequences unique to the SK clade. This script was used between SK_AV1 strain and non-SK strains with a minimum size of 5,000 bp. RODs were then BLASTed against each other (60% coverage and 60% identity cut-off) to determine their presence in the other genomes. Finally, RODs, phages, and GIs where visualized in BRIG [56], using the strain SK_AV1 as reference genome for the alignment and all their genes were re-annotated with WebMGA (e-value of 10^−5^ [57]).

### Genome degradation

DFAST [58] was used to determine pseudogenization resulting from frameshift mutation and nonsense-mutation in the genes of the 17 *C. maltaromaticum* genomes using the LMA 28 strain as a reference. Pseudogenes where annotated with WebMGA [57].

## Results

### Strains identification

First, we investigated the phyletic affiliation of the nine *C. maltaromaticum* SK strains isolated from the infected brain and inner ear of 5 stranded sharks using phylogenetic and phylogenomic approaches (Table 1). All the nine strains were identified as *C. maltaromaticum* according to complete 16SrRNA gene analysis (Fig 1A), the Tetra-nucleotide correlation analysis (TCS >0.99), and the Average Nucleotide Identity (ANIb >98%). Furthermore, 16S rRNA sequences from all publicly accessible genomes of *Carnobacterium* (n=37) (S2 Table) revealed little phylogenetic divergence among sequenced genomes of *C. maltaromaticum* with the exception of the *C. maltaromaticum* 757 CMAL, derived from human skin (Fig 1A). However, both the identification of 91,236 identified SNPs distributed throughout the core genomes of the 17 *C. maltaromaticum* genomes (Fig 1B) and the clustering based on functional genes (FigFams) distribution (S1 Fig), produced a more distantly branched cluster with the nine SK strains, suggesting the shared evolution of the SK strains and their divergence from other known isolates. The 17 *C. maltaromaticum* strains were classified into three groups, diseased-shark (n=9), diseased-trout (n=2), and food-associated isolates (n=6).

**Fig 1.**
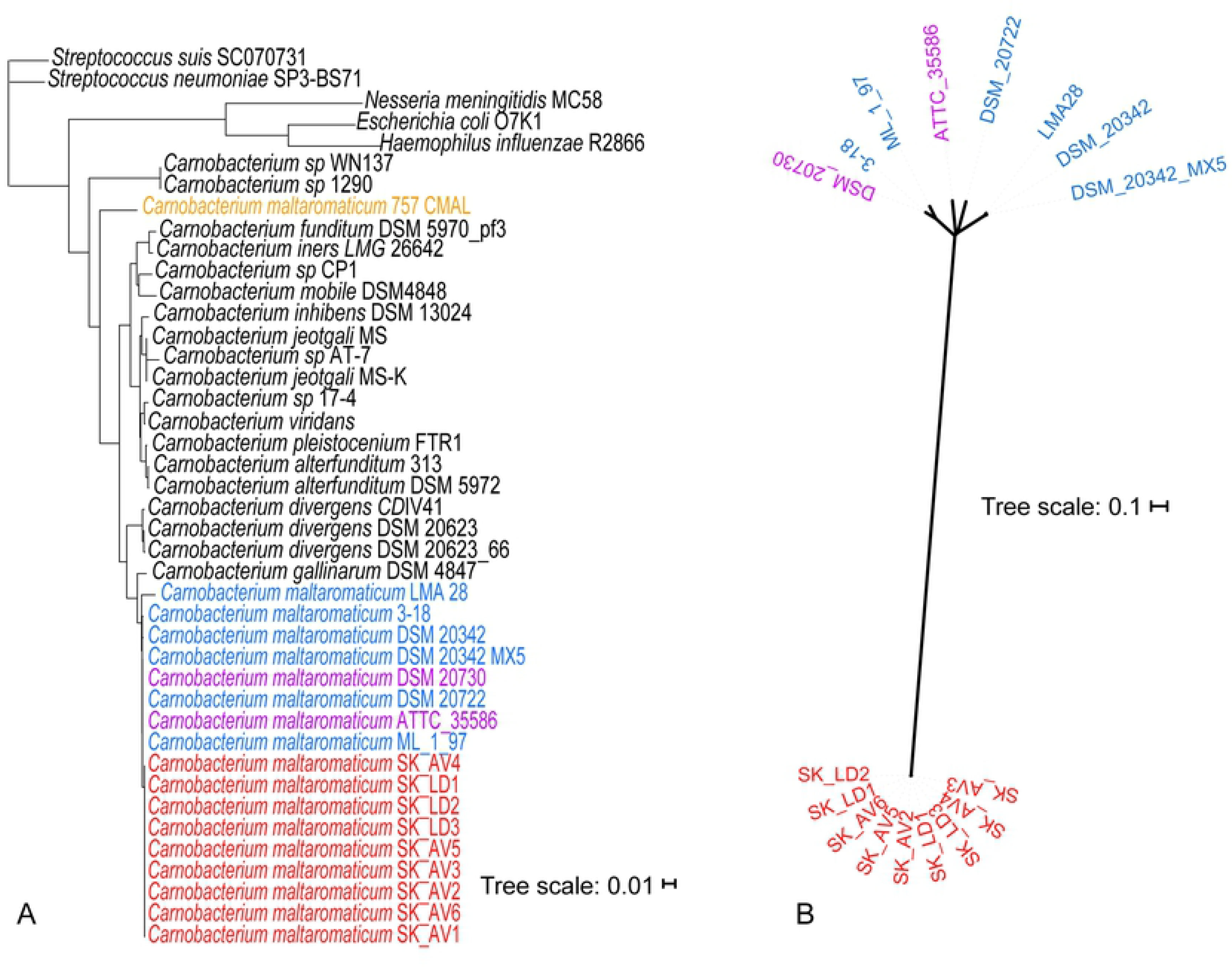
*C. maltaromaticum* phylogeny according to the 16S rRNA gene and SNPs. (**A**) Neighbor-joining tree of the 16S rRNA gene of 37 *Carnobacterium sp.* along with 5 pathogenic non-*Carnobacterium* strains as outgroup. Scale represents the frequency of substitution per site. (**B**) Unrooted neighbor-joining tree according to the SNPs in the core genome of the 17 C. maltaromaticum strains. Isolates from diseased-sharks: red, diseased-trout: purple, food: blue, and human skin: orange.

### Genomic plasticity & Gene content

Considering the 17 *C. maltaromaticum* genomes (Table 1), the pan-genome neared completion (Heaps law model, α = 0.93, S2 Fig) and consisted of 4,746 gene clusters (3,074.6 ± 136.9 clusters per genome) annotated with 1,444 different functions (Table S3). Conversely, the binomial mixture model, anticipated a final core and pan-genome size of 2,285 and 5,903 gene-clusters, respectively, suggesting potential missing gene clusters from the *C. maltaromaticum* pan-genome. As shown by the actual gene distribution among the *C. maltaromaticum* (Fig 2), there was relatively high gene diversity among the 17 genomes, reflecting their diverse life-styles and isolation sources. However, most of the genes in the SK strains belonged to the SK-clade’s core-genome, with the accessory genes representing a smaller proportion of the genome (Fig 2). Thus, most of the genome diversity observed within the *C. maltaromaticum* group reflects variation in the gene content outside the SK clade.

**Fig 2.**
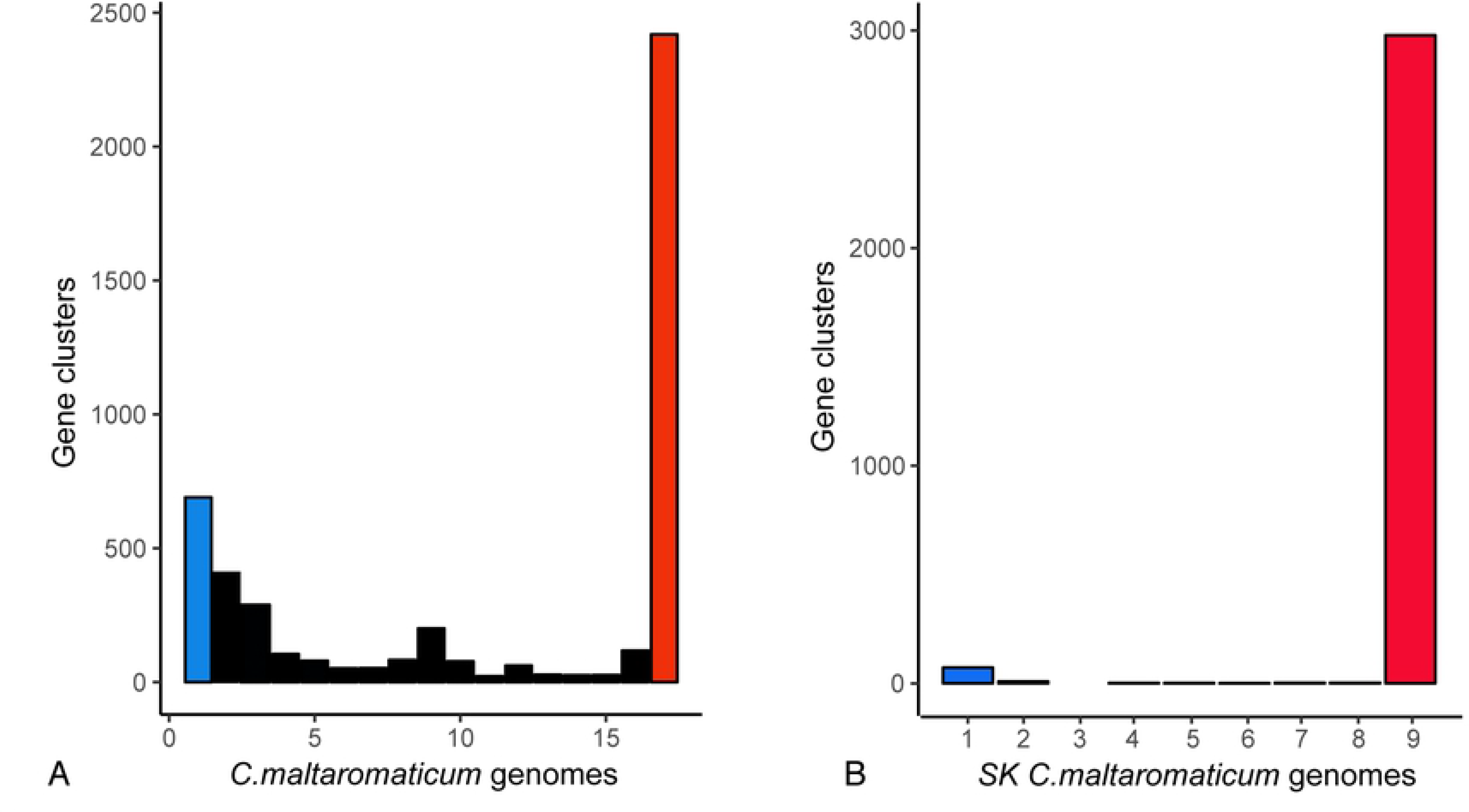
Core, accessory and singleton genes within the *C. maltaromaticum* pan-genome. (A) Distribution of gene clusters in the 17 *C. maltaromaticum* genomes. (B) Distribution of gene clusters in the 9 *C. maltaromaticum* SK genomes. Bar plot shows core genome (red), singletons (blue) and accessory gene clusters found in more than 1 and < 17 genomes (black bars). Comparison of both Figs reveals that SK strains have few genes different from each other while the accessory genome in all 17 strains represents a big proportion of the pan-genome.

The core, accessory, and singleton genes in the 17 strains represented 50.9%, 34.5% and 14.5% of the identified genes, respectively (Fig 2). In addition, 33.1% of the identified gene clusters, mostly in the accessory and singleton category, remained with unknown function. Only 106 clusters unique to the SK strains had functional annotation. Among these, only 19 COG functions were not shared by the other *C. maltaromaticum* genomes (Table S4), and only 9 were present in all the SK strains. Apart from the *unknown functions*, predominant COG functions in the core genome were associated to housekeeping genes such as *Translation, ribosomal structure and biogenesis* (12.35 %)*, Amino acid transport and metabolism* (9.35 %). Conversely, the accessory genes and singletons, involved in the adaptation to the environment, had a higher occurrence of functions involved in *Mobilome: prophages, transposons* (accessory 12.13 %), *Cell wall/membrane/envelope biogenesis* (singletons 11.11%), *Defense mechanisms* (singletons 11.11 %), *Extracellular structures* (singletons 8.33%) and *Cell motility* (singletons 8.33%) (S3 Fig).

The systematic comparison of the 17 *C. maltaromaticum* genomes highlighted 6 GIs conserved in all the SK genomes, four of them being unique to the SK clade (Fig 3). A complete phage sequence (i.e., PHAGE_Strept_phiARI0746_NC_031907 from *Streptococcus pneumoniae* 10B04751, Phage 4) was also identified. Finally, two incomplete phages (Phages 2 and 3) and one potential phage (Phage 1) sequences were also found in all SK genomes (Fig 3). However, no virulent gene was found in the phages or GIs when aligning the corresponding sequences against the VFDB. In total 35 RODs were identified, but only 5 were specific of the diseased-shark strains and 4 were also shared by at least 1 of the diseased-trout strains (Fig 3). Although most of the genes identified in the phages, GIs, and RODs had unknown functions (70%, 71%, and 79%), many genes with annotated function in the RODs were associated with cell wall synthesis and capsular polysaccharide production (S5 Table).

**Fig 3.**
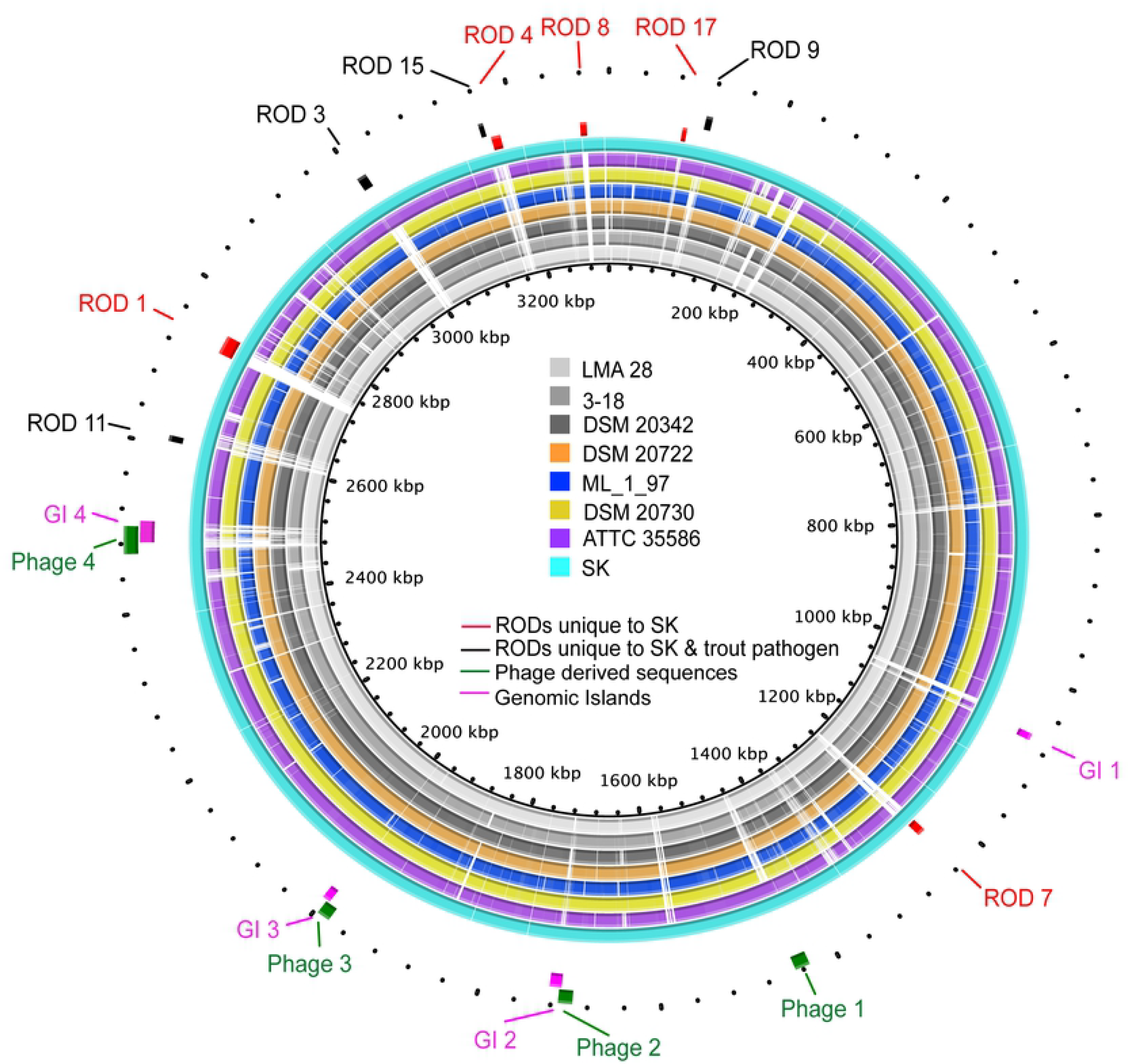
Alignment of *C. maltaromaticum* genomes against SK AV1 strain. The outer light blue ring contains all the SK clade (besides the AV1, which was used as a reference for the alignment). RODs in red are unique to shark and trout pathogen, whereas the black RODs are unique to the SK clade. Green represent phage-derived sequences identified by PHASTER and pink represent GIs unique to the SK clade identified by IslandViewer.

### Functional variation between groups

When comparing the distribution of predicted functions in the analyzed genomes, we identified significant differences between the SK strains (n=9 genomes) and the food-associated strains (n=6 genomes) whereas no significant difference was observed when considering diseased-trout strains (n=2 genomes, data not shown). More precisely, the SK strains had significantly less genes for *Replication, recombination, and repair* and *Mobilome: prophages, transposons*. In addition, the SK strains consistently displayed reduced number of genes for *Cell motility* although not significantly. This mirrored the variable number of genes for this function in the non-SK strains. Conversely, the SK strains were significantly enriched in genes for *Secondary metabolites biosynthesis, transport and catabolism, Post-translational modification*, and *Energy production and conversion*, among others (Fig 4).

**Fig 4.**
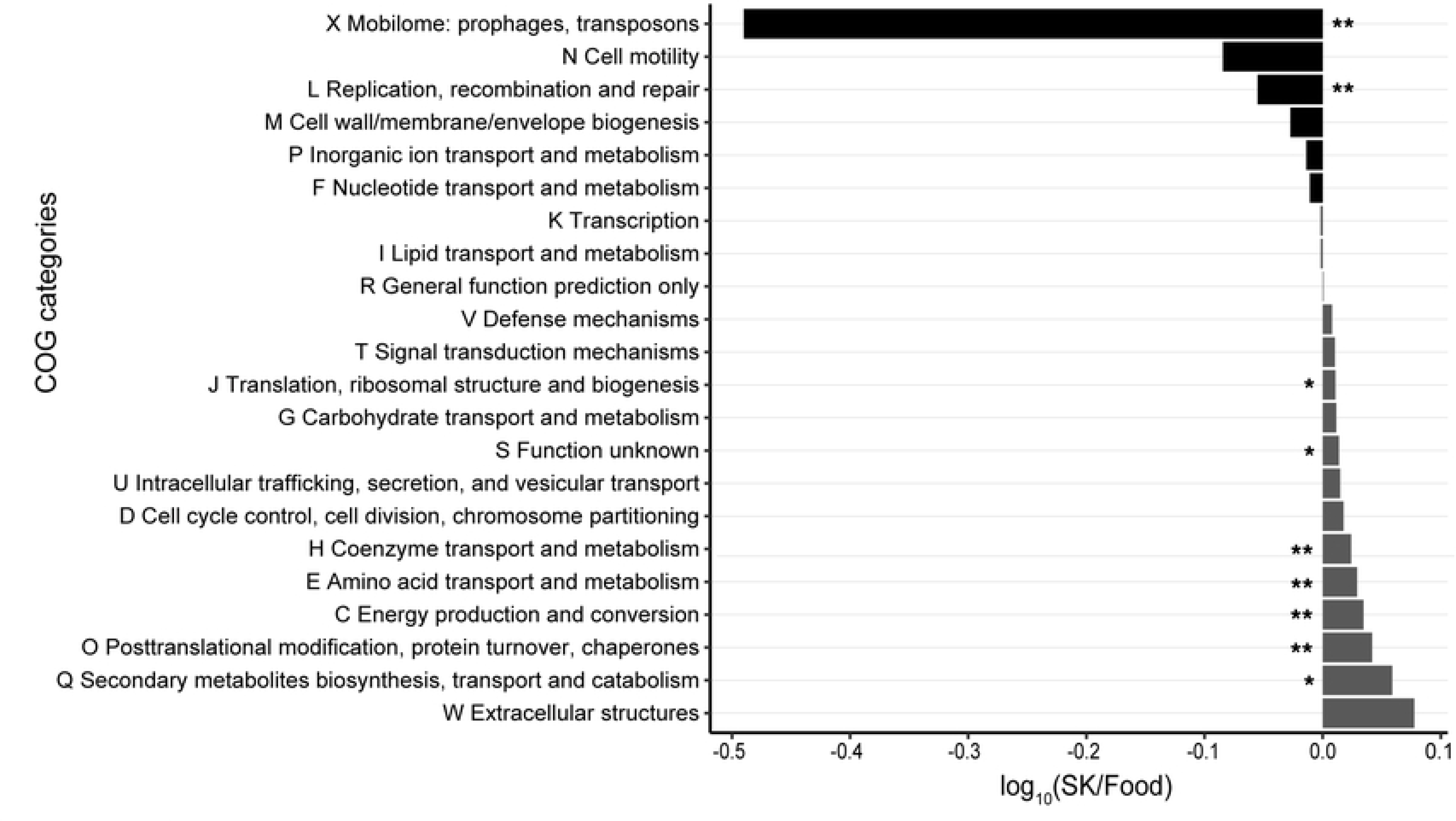
Functional comparison between strains isolated from food and diseased sharks. X axis represents the log ratio of the COG categories relative abundance. Asterisks indicate level of significant (one asterisk p < 0.05, two asterisk p < 0.01) when performing a Wilcoxson Rank Sum test with Bonferroni correction for multiple comparison. Variations between these groups indicate their adaptation to different environments associated different functional needs.

The SK strains and the other *C. maltaromaticum* genomes contained 29.3 ± 1 and 29.3 ± 4 virulent associated genes, respectively. However, seven were only found in the SK strains. These genes coded for a potential enzyme involved in capsular polysaccharide synthesis, a dTDP-4-dehydrorhamnose reductase, and a prolipoprotein diacylglyceryl transferase all found in the 9 SK strains. In addition, a PixD protein, a bifunctional aldehyde-alcohol dehydrogenase, a cytotoxic necrotizing factor 1, and a fimbriae were identified only in some of the SK strains (S6 Table). Using the FigFam annotations, we compared and identified unique functions inside the *Virulence, Disease and Defense* category (S4 Fig). SK strains had an overrepresentation of genes for the *Invasion and intracellular resistance* (with a SK-unique internalin like protein *Lmo0327*), and *Bacitracin* (with a SK-unique bacitracin stress response regulator). Furthermore, the SK clade showed unique presence of fosfomycin resistance and beta-lactamase along with other genes involved in the synthesis of cell wall and capsule, phages, iron metabolism, RNA and nucleotide metabolism, respiration and stress response (S7 Table).

### Pseudogenization

Within the 17 *C. maltaromaticum* genomes, the SK strains displayed a high frequency of pseudogenes (263 ± 1) resulting mostly from non-sense mutations (Table 1). Exceptionally, the *C. maltaromaticum* ML_1_97, isolated from fresh salmon, also had a high number of pseudogenes, perhaps as a consequence of the high number of contigs of this genome assembly (Table 1). Pseudogene functional annotation in SK strains showed high frequency of genes annotated under the categories of *Carbohydrate transport and metabolism* (13.1%), *Amino acid transport and metabolism* (10.6%), and *Energy production and conversion* (9.7%) (S5 Fig). Pseudogenization potentially inactivated 115 functions in the SK strains and caused the strains from food and diseased-trout to have a total of 166 and 126 functions absent from the SK cluster, respectively.

## Discussion

*Carnobacterium* are environmental microbes [6] frequently identified in dairy products [59], whereas some species are able to cause disease in fishes [11, 14, 15]. Members of this genus are hard to identify and discriminate using standard techniques (e.g., phenotypic characterization, 16S rRNA phylotyping) because of their phylogenetic proximity and functional plasticity [11, 60]. However, a recent study using Multilocus Sequence Typing revealed that *C. maltaromaticum* causing disease in teleosts are non-clonal and vary from isolates from dairy products [61].

Recently, *C. maltaromaticum* isolates have been isolated from the brain and inner ear of stranded salmon sharks [1] and common thresher sharks [2] found with severe brain and inner ear infection. To date, both the detailed epidemiology and the exact etiology of these infections remain largely unknown; however, the consistent presence of *C. maltaromaticum* in infected areas highly suggests its role in the development of the pathology, as it is involved in the development of disease in bony fishes [11, 14, 15]. Here, we performed a thorough comparison of *C. maltaromaticum* sequenced genomes in order to identify the genomic features associated with niche adaptation in this clade. As expected, the systematic comparison of sequenced genomes from the *Carnobacterium* genus highlights the phylogenetic cohesion of all the *C. maltaromaticum* isolates displaying less than 1% 16S rRNA variation [11, 60]. The phylogenetically coherent *C. maltaromaticum* cluster, containing the SK strains, is related with the *C. divergens* and *C. gallinarum* species [59], two species frequently detected across environments [6], whereas the other species formed a distinct and more distantly related cluster. Interestingly, the clustering of *C. maltaromaticum* SK strains according to phylogenomic markers (e.g., SNPs, functional gene distribution) correlated with their phylogenetic clustering [44]. Thus, all the SK isolates are distantly related to other lineages of *C. maltaromaticum* used in this study, and share a unique specific gene repertoire. This suggests that SK strains share the same evolutionary history and do not seem to originate from food or teleost contamination sources [12, 13].

Since the genomic plasticity of bacteria underlines the ability of phylogenetically related microbes to colonize distinct environments [62] and sometimes to be involved in pathologies [19, 21, 63], we expected the SK strains to display specific genomic adaptation of the host-adapted lifestyle. Frequently identified features associated with this life-style include genomic reduction mirrored by the reduced genome and pseudogenization [24, 25]. Furthermore, the presence of specific genes, such as the virulence genes, can aid in the invasion process [52]. Genome streamlining is a gradual process, which erodes genes and functions potentially supported by the host, and eventually results in a reduced genome with only essential functions conserved. Similar genome reduction is observed in well-known pathogens *Shigella* sp. and *Salmonella enterica* [64, 65]. Many of the pseudogenes identified in the SK strains most likely affected the functionality of genes as a consequence of the introduction frameshift, indels, and random stop codons in the coding regions (Table 1). The accumulation of pseudogenes in sequences likely results in the loss of function, suggesting a stronger connection between the microbe and its host [25, 66]. Here, the pseudogenization of SK genomes affected mostly genes involved in central metabolism (e.g., *Amino acid transport and metabolism*). Similar pseudogenization has been identified in pathogens with the host’s cellular machinery supporting the loss of function [67, 68]. However, as genomic degradation reduces the fitness in the open environment, losing too many functions or of essential functions could have a detrimental effect on the ability of the strain to propagate [69, 70]. Thus, unlike some highly specialized pathogens, many diseased-causing agents rely on their ability, even reduced, to survive and potentially propagate in the environment [71].

Interestingly, apart from the described loss of accessory genes and the pseudogenization of some important functions, the SK strains displayed a specific set of intact genes potentially associated with the development of the pathology. Among others, SK strains have two extra genes encoding ABC transporter for polysaccharide export and potentially involved in exopolysaccharide production that could promote the evasion of the immune system of the shark [72–74]. In addition, *CapO* (S6 Table), a gene essential in the for capsule synthesis [75], is only found in SK strains. Furthermore, some peroxiredoxin genes, potentially involved in protecting the bacteria from oxidative stress mediated by the host immune system [76–78], were also more abundant in the SK strains (S6 Table). Finally, an internalin-gene, supporting the colonization of the host by facilitating the crossing of the intestinal epithelium and blood-brain barrier [79] was also unique to the SK strains (i.e., *Lmo0327*) (S6 Table).

The low frequency of genes from Mobile Genetic Elements in the SK strains relative to the other food-derived and diseased teleost-derived genomes (Fig 4), could be due their limited interaction with environmental microbial communities, which limits the potential to gain new functions through horizontal gene transfer and further reduce the genome diversity. This phenomenon is often found in pathogens [80]. Nonetheless, a complete phage and four GIs (three of likely phage origin) were specific to the SK clade (Fig 3). Additionally, the SK strains displayed nine RODs of unknown origin, with four being shared with the trout pathogen (i.e., *C. maltaromaticum* ATTC 35586). The phages and the RODs sequences, although containing many genes with unknown function, displayed several genes involved in cell-wall and exopolysaccharide biosynthesis.

In total, we compared 17 *C. maltaromaticum* bacterial genomes in order to identify the genomic features associated with the 9 strains isolated from diseased-sharks. However, we recognize that many of these genomes are not complete and that the nature of the missing information could affect our results and interpretation in different ways, for example by inferring the wrong number of genes (potentially adding or subtracting genes due to the fragmentation of the genome into multiple contigs, and the presence of gaps) [81–83]. However, most of the genomes used in this study had low contig number (19.9±5 for SK strains and 34.9±28 for remaining seven, with the exception of ML 1 97 which had 176 contigs), and large contig size (N50 > 312,000 for SK strains and N50 > 75,000, for all genomes with the exception of strain ML 1 97 with N50 = 26,642) (Table 1), which decrease the potential for missing or false produced genes [81]. Nonetheless, over or under-representation of gene numbers could be affecting the identification of specific genomic features. However, sequencing 9 SK genomes confirmed the presence of features of interest in all strains and reduced the possibility of missing genes. Additionally, for consistency, all the publicly accessible genomes included in this study were re-annotated and treated in the same way in order to minimize systemic bias introduced by different bioinformatics pipelines.

Contrary to high diversity among teleost pathogens [61], the strains isolated from these diseased-sharks have very similar genomes, with only few identified accessory genes. The conservatism of identified genomic features such as the functional genes and the pseudogenization in the SK strains supports a monophyletic origin for the nine SK strains rather than a converging evolution. Indeed, the nine SK strains, despite being derived from different tissues (brain and inner ear), and sharks (i.e., common thresher vs. salmon shark), collected in different years (from 2013 to 2016) and locations (throughout California) (S1 Table), are all highly similar. Thus, the consistent presence of *C. maltaromaticum* in diseased sharks, and the proximity among the SK strains (Fig 1B) further suggest that *C. maltaromaticum* SK strains are involved in periodic thresher and salmon shark stranding events.

Bacterial pathogens have a range of possible classifications from environmental and commensal organisms that occasionally cause infection to obligate pathogens not existing outside of their host [63]. According to the SK genomic profiles and its consistent association with shark disease, we can hypothesize this clade as opportunistic pathogen, as it likely maintains the capability to survive and propagate while outside the host. This would benefit the spread of *C. maltarmaticum*, as host-to-host transmission seems unfeasible in these species of shark, which do not interact with each other apart from mating and during gestation periods. While few sharks strand every year, the exact proportion of the populations affected by the disease is unknown, as only those that make it to the beach are accessible. However, at least two species of shark seem to be affected by *C. maltaromaticum* SK strains, thus suggesting other sharks could possibly host and be affected by some of these strains. To further our knowledge on the environmental source of this bacterium and its implication in the development of the pathology, future studies should focus on determining the presence of SK strains in different environmental settings and hosts. In this context, the identification of conserved SK-specific genomic regions will help track SK-isolates in sequenced microbiomes. Additionally, using those unique regions, SK-specific sets of primers will help identify SK strains in heathy/diseased sharks and their environment, identify the geographic distribution, the host range, and the mechanistic implication of SK strains in shark stranding.

## Supporting information

**S1 Fig. Complete-linkage clustering of 41 *Carnobacterium sp*. and 5 outgroups.**

Clustering was performed from a Bray Curtis dissimilarity matrix according to FigFam annotation. *C. maltaromaticum* strains derive from diseased sharks (red), diseased trout (purple), food (blue) and human skin (orange).

**S2 Fig. *C. maltaromaticum* pan-genome accumulation curve.**

Curve shows the increment of unique gene clusters with the addition of new genomes to the pan-genome analysis. Blue area represents confidence intervals from standard deviation.

**S3 Fig. Functional distribution of the core, accessory and unique genes of the *C. maltaromaticum* pan-genome.**

Gene clustering and annotation are according to Anvi’o. Plot shows that high proportion of the core genome are house-keeping genes (like metabolism), while non-essential genes represent a higher proportion of the variable genome indicating difference in adaptations between strains.

**S4 Fig. Scaled heat map of the genes categorized as virulence by the RAST annotation server in all *C. maltaromaticum*.**

SK strains highlighted in bold show higher proportion of genes that represent invasion and resistance, perhaps indicating their tendency towards a more pathogenic lifestyle.

**S5 Fig. Functional gene distribution of pseudogenes extracted from the SK genomes.** Pseudogenes were predicted according to COG annotation using WebMGA software. High proportion of the genes are associated with metabolism, indicating the potential loss of metabolic functions that are no longer essential when pathogens adapt towards an intra-cellular lifestyle.

**S1 Table. *C. maltaromaticum* strains isolated from diseased sharks.** The column “Sharks” indicates the shark species and region the strain was isolated from. Numbers following shark species name used to discriminate between different sharks sampled. Column “Location” indicates where the shark was found stranded.

**S2 Table. Strains used in the phylogenetic analysis.**

**S3 Table. Pan-genomic distribution of the *C. maltaromaticum* strains.** Table shows total number and percentage of gene clusters forming the pan-genome, core genome, accessory genome and singletons. Number of clusters and relative abundance of annotated and unclassified genes are also shown along with the number of different functions in each category. The pan-genome of just the SK clade is shown separately with their gene annotation and functional distribution.

**S4 Table. Unique COG functions to the SK clade.** The column number of genomes represent presence of the function in just one or all SK genomes.

**S5 Table**. **Assigned functions to the genes found in the RODs.** Table shows genes in RODs unique to SK strains and unique to SK and trout pathogen genomes. Table reveals all genes with known function, while most of the genes were unclassified (not shown here).

**S6 Table. Virulent associated genes found in all *C. maltaromaticum* genomes according to the VFDB.**

**S7 Table. Unique functions to the SK clade according to RAST FigFam-based annotation.**

